# First report of *Hexamita nelsoni* in blue mussel (*Mytilus edulis*): morphology, phylogeny, and host-parasite interaction

**DOI:** 10.1101/2024.09.18.613806

**Authors:** Anders Alfjorden, Jon Jerlström Hultqvist, Staffan Svärd, Fabien Burki

## Abstract

Bivalve diseases caused by protozoan parasitic infection are monitored by coordinated surveillance programs across European member countries. These routine monitorings of bivalve health, however, only survey a few relatively well known parasites, which may leave undetected a range of lesser known opportunistic protozoan agents. Under increased stress, for example due to shifting environmental conditions affecting water quality and nutrient access, these opportunistic parasites may develop pathogenicity impacting reproduction or growth and therefore decreasing the success and future development of aquaculture or wild population sustainability. During routine histopathological surveillance of blue mussels in Sweden, we noticed cryptic lesions in the adductor muscle in high prevalence over a period ranging from 2017 to 2022. These lesions suggested cell infiltration and degenerative changes in the muscle tissues. In this study, we report on the isolation, identification, and culture of the protistan cells likely responsible for at least some of these lesions. Using histology and cytology, molecular phylogeny, and catalyzed reporter deposition fluorescence in situ hybridization (CARD-FISH), we show that the cells correspond to the *H. nelsoni* sequence initially isolated from oysters. We also reveal the presence of *H. nelsoni* in the digestive glands and mantel, and that this parasite might be horizontally transmitted from adult to larvae via infections within gonads, ovaries. This study represents the first report of *H. nelsoni* in blue mussel.

## 1. Introduction

Bivalves are filter feeders that continuously consume microorganisms, including single-celled protists. Some protists can survive within the host’s filtration and intestinal systems, avoiding host defense mechanisms and feeding on host resources or other components of the microbiome, and occasionally turning parasitic and/or pathogenic (Bass and del Campo, 2020; Carballal et al., 2001; Gestal et al., 2008; Villalba et al., 1997). Known pathogens are routinely monitored by national and international targeted surveillance programs such as the EU national reference laboratories (NRL) monitoring programs, run by individual member countries according to the EU animal health law (EUR-Lex, regulation 2016/429). In these programs, representatives from EU countries meet yearly to report from their activities to mitigate mass-mortality events affecting aquaculture as well as wild stocks. Notably, these programs notify detection of different pathogens, creating notification lists, biogeographic alerts and evaluating the risk for disease (Fox et al., 2020) as well as establishing legislative frameworks (Bass et al., 2015; Carnegie et al., 2016; Carnegie and Engelsma, 2014; Hine and Zealand, 2002; Webb, 2008). Despite these routine surveillance programs, bivalves are also likely hosts to an array of opportunistic parasites that are not monitored but can, under limiting conditions, negatively impact reproduction or growth and therefore decreasing the success and future development of bivalve aquaculture and the sustainability of wild stock (Bass et al., 2019; Bass and del Campo, 2020). Emerging or unlisted pathogens can thus be missed by targeted molecular diagnostic tools, an issue made worse by the decreasing use of broad spectrum microscopic observations (Carnegie et al., 2016). Climate change has been shown to be an important inducing factor of opportunistic infections and with a changing climate it is expected that the problem with opportunistic infections will increase in aquaculture (Collins et al., 2020).

Several decades ago, studies of oyster hatcheries and aquaculture farms reported that flagellated cells belonging to the genus *Hexamita* (Metamonada, Diplomonadida, Hexamitinae; (Kolisko et al., 2008)) were associated with winter mortalities in relation to poor husbandry or bad water quality (Elston, 1994; Mackin et al., 1951). These studies confirmed the presence of *Hexamita* sp. in bivalves, which were first detected in the stomach of an oyster in (Certes, 1882). Although the nature of the relationship of *Hexamita* to the oyster hosts has remained ambiguous (Feng and Stauber, 1968; Mackin et al., 1951; Scheltema, 1962; Schlicht and Mackin, 1968; Vivares et al., 1987a, 1987b), most known diplomonads live as commensals or parasites of animals, including the human parasite *Giardia intestinalis* (Ankarklev et al., 2010) and the fish parasites of the genus *Spironucleus* (Xu et al., 2014). In addition to parasites, several free-living diplomonads mostly from freshwater samples were recently described by detailed ultrastructure and phylogenetic analyses (Mazancová et al., 2023). Taken together, our current limited knowledge of diplomonads, suggests a larger diversity of free-living lineages than previously understood, which seem to have independently evolved from endobiotic ancestors several times (Kolisko et al., 2008; Mazancová et al., 2023).

To date, only two *Hexamita* species have been described, the free-living *H. inflata* and *H. nelsoni* found in oysters (Schlicht and Mackin, 1968; Vivares et al., 1987b). The phylogenetic placement of several species initially proposed to be related to *Hexamita* isolated from various animals such as birds, mammals, and fish (Ferguson, 1979; Griffiths, 1971; Hussain, 2001; Poppe et al., 1992) have been revised by morphology, ultrastructure and molecular data and often been transferred to the genus *Spironucleus* (Brugerolle et al., 1980; Ferguson, 1979; Jørgensen and Sterud, 2007; Sterud et al., 1998). The recent study of free-living diplomonads has further shown that *Hexamita* is a polyphyletic genus and genetically split into at least four different clades (Mazancová et al., 2023). Thus far, *Hexamita* has not been reported from blue mussels (*Mytilus edulis*).

During routine histopathological surveillance of blue mussels in Sweden, we noticed cryptic lesions in the adductor muscle in high prevalence over several years from 2017 to 2022. These lesions suggested cell infiltration and degenerative changes in the tissue. In this study, we report on the isolation, identification, and culture of the cells likely responsible for at least some of these lesions. Using histology and cytology, molecular phylogeny, and catalyzed reporter deposition fluorescence in situ hybridization (CARD-FISH), we show that the cells correspond to the *H. nelsoni* sequence initially isolated from oysters. We also reveal the presence of *H. nelsoni* in the digestive glands and mantel, and that this parasite might be horizontally transmitted from adult to larvae via infections within gonad ovaries. This study represents the first report of *H. nelsoni* in blue mussel.

## 2. Material and methods

### 2.1. National surveillance and sampling of blue mussels

Wild and farmed blue mussels were collected yearly during the Swedish national monitoring program for marteilioses (*Marteilia spp*.) between 2017 and 2022. 30 specimens were collected in each of 5 different sampling areas at the Swedish westcoast and investigated by histology (starting from Värobacka in south and ending at Strömstad, close to Norway in the north). This program follows the EURL mollusc screening recommendations, using approved guidelines for the investigation of notifiable diseases within the European Union (Aquatic animal health law, EUR-Lex, regulation 2016/429). Histology is the general method for mollusc diagnostics to find notifiable parasitic diseases in bivalve hosts. However, this method can also reveal new or emerging diseases when sudden and unexpected mortalities occur.

### 2.2. Cell isolation, live cell imaging and sample processing

Biopsies (∼5×5×5mm) taken from the posterior adductor muscle were inoculated in Eagle’s Minimal Essential Medium (EMEM) supplemented with calf serum and antibiotics, normally used for fish virus surveillance purposes (https://www.eurl-fish-crustacean.eu/fish/diagnostic-manuals). In this study, the culture medium was modified to a higher salinity (∼2%). This medium, hereby named Marine EMEM, is prepared by use of EMEM supplemented with serum and antibiotics (Penicillin/Streptomycin) mixed at a ration of 1:1 with sterile (autoclaved) filtered seawater or Red Sea salt dissolved to a concentration of 3% NaCl. The muscle biopsies were placed in top-filled “slanted Leyton tubes” (NUNC^TM^ product nr. 145470) and kept at low temperature (2-6°C). The culture medium was exchanged within the first 12 hours and once again within one week but then kept for two months and regularly monitored by light microscopy (inverted microscope, Nikon TMS). When flagellated trophozoites were growing to high densities, subsamples were taken to purify the culture by transfering to new slanted tubes with marine-EMEM. Bottom cultures were re-isolated using Pasteur pipette, externally washed in 70% ethanol for surface sterilization.

Isolates of live trophozoites were inspected by phase contrast (PH) using Nikon Labophot, Tokyo, and microscope lense, 100/1.25 oil 160/0.17 Ph4. Cells were documented by a Canon EOS 500D attached to the microscope by a lens-adaptor (Martin Microscope Company MM-SLR adaptor S/N:0468; Easley, SC USA). The cell-cultures were also investigated by an inverted microscope Nikon eclipse Ts2R, using Nikon 100x/1.25 oil PH3 DL and also inspected by differential interference contrast (DIC) using the Nikon eclipse with Plan Fluor DIC, 100X/1.30 oil lens. Images were documented using the attached TIS Camera USB 3.0 Monochrome 1/1.2 CMOS.

### 2.3. PCR, cloning, sequencing and phylogenomics

The first isolates of flagellated trophozoites (M1-4652-18 and M29-2652-18) were pelleted by centrifugation (2000g x 2 min) using 1.5 ml aliquots of culture followed by discarding the supernatant. The collected cells were digested using Proteinase K and ATL buffer at 56°C overnight, under slow shaking condition (60 rpm). DNA was isolated using Qiagen DNeasy Blood and tissue kit, following the manufacturer’s instructions. The eluted DNA concentration was measured on a Qubit Fluorometric Quantification Machine (Thermo Fisher Scientific).

The 18S rDNA gene was amplified using a combination of broad ranging eukaryotic primers: PF1 5′-TGCGCTACCTGGTTGATCCTGCC-3′ (Keeling, 2002) and FAD4 5′-TGATCCTTCTGCAGGTTCACCTAC-3′ (Deane et al., 1998; Medlin et al., 1988). The PCR reaction was conducted using EconoTaq PLUS 2× Master mix (Cat no. 30035-1, Lucigen, LGC Biosearch technologies, USA) as follows: Initiation, 94°C for 2 min; 35 cycles of 95°C for 30 s, 50°C for 1 min, 72°C for 1.5 min and final extension, 72°C for 10 min. PCR products were cloned using StrataClone PCR cloning kit (product no: 240205, Agilent Technologies Sweden AB, Kista) and reamplified with M13F/M13R primers using DreamtTaq (Thermo Fisher Scientific, Baltics UAB, Vilnius, Lithuania). Bands of the expected size were purified with Illustra^TM^ ExoProStar^TM^ 1-step Enzymatic PCR and Sequence reaction clean up Kit (prod no. US77702, GE Health Care UK Limited, Buckinghamshire). Purified clones were Sanger-sequenced in both directions at Macrogen (Macrogen Europe BV, Amsterdam, Netherlands). The sequences were verified, trimmed, and assembled with Geneious v.10.0.9 (Kearse et al., 2012)

One of the new isolates (M2-6738-22) was sequenced more extensively with Nanopore to retrieve the whole ribosomal operon (18S rDNA to 28S rDNA). This isolate was also later used for more detailed microscopic investigations (see below). Trophozoites (400 ml of culture) were enriched by centrifugation for 10 min, 500×g, 4°C. The cells were resuspended in 1 ml culture supernatant filtered through 0.2 µm filter (CSF) to remove larger biofilm fragments. The cell suspension was centrifuged on a 9 ml Histopaque gradient and the cells were recovered from the Histopaque-CSF interface by pipetting. The recovered cells were diluted in 10 ml CSF and repelleted by centrifugation as described earlier. The cell pellet was lysed in 2 ml lysis buffer (1% SDS, 50 mM EDTA, 30 mM DTT and 800 µg Proteinase K (Thermo-Scientific, EO0491) at 56°C for 2 hrs. Nucleic acid was extracted using two rounds of Phenol-Chloroform-Isoamylalcohol (25:24:1) and one round of Chloroform-Isoamylalcohol (24:1) separation followed by precipitation using 0.7× volumes of isopropanol. The isolated nucleic acid was treated with 500 µg of RNase A for 30 min at 37°C. The DNA was isolated using a Qiagen Genomic Tip G/20 according to the manufacturer’s instructions. DNA concentrations were determined using the Qubit dsDNA BR kit using a Qubit 2.0 (Invitrogen). The size distribution of the DNA was determined using agarose gel electrophoresis. DNA purity was determined by NanoDrop 1000 Spectrophotometer v3.8.1 (Thermo Scientific).

Library preparation was accomplished using 1500 ng genomic DNA using the Oxford Nanopore ligation sequencing gDNA kit (SQK-LSK112), version GDE_9141_v112_revE_01Dec2021 and sequenced on an R10.4 flow cell (FLO-MIN112) using MinION Mk1C running MinKNOW v21.11.6 software. The raw nanopore data were basecalled using Dorado version 0.4.0 in super-accuracy duplex mode. Adapter removal for Nanopore was done using Porechop v0.2.3. (Wick et al., 2017). The Nanopore reads were filtered using Filtlong v0.2.1 (https://github.com/rrwick/Filtlong) to remove reads less than 1000 bp and the worst 5% of reads. The Nanopore datasets were assembled using Flye v2.9-b1768 (Kolmogorov et al., 2020)) with the --nano-hq, --meta flags and otherwise default settings. A contig (contig_1143) containing the Hexamita ribosomal loci was identified in the Flye assembly by BLASTN using the 18S rDNA sequence from *Hexamita nelsoni* (GenBank ID: EF050053) as the query. The Hexamita 18S, 5S, and 28S rDNA gene sequences were identified by searching the contig at RNACentral. The *Hexamita inflata* 18S, 5S and 28S rDNA were identified by the same procedure in the assembly described by (Akdeniz et al., 2024) (GenBank assembly: GCA_963988835.2).

For phylogenetic inferrences, a broad diversity of Hexamitinae sequences were selected based on available studies of aquatic diplomonad diversity (Jørgensen, 2007; Kolisko et al., 2008; Mazancová et al., 2023) and downloaded from GenBank. A list of all sequences used in this study is presented in Supplementary Table 1. These sequences were then investigated together with our cloned sequences from the blue mussel isolates to phylogenetically place the *Hexamita* spp. flagellates. All isolates from the Swedish blue mussels are listed in Supplementary table 2, together with *Hexamita inflata* isolated from shallow water in a wetland in Řevnice, Czech Republic. We used IQ Tree multicore version 2.2.2.6. (Minh et al., 2020) to infer a maximum likelihood (ML) tree, after investigation by the modelfinder algoritm (Kalyaanamoorthy et al., 2017) for the best fitting nucleotide substitutions model: GTR+F+R6. The result from this phylogenetic investigation is presented in Figure 3. The sequences were aligned with MAFFT v7.490 (Rozewicki et al., 2019) using the mafft-linsi algoritm. TrimAl v. 1.2rev59 with gap treshold settings: 0.01 and string treshold 0.001 where used to trim unambigous sequnces. The branch support was assessed with standard non-parametric bootstrap based on 100 bootstrap replicates.

### 2.4. Histopathology and cytology

Samples were collected as cross-sections of the mussel, covering organs such as gills, mantel, digestive glands and foot, according to EU guidelines (https://www.eurl-mollusc.eu) but also in line with the dissection description for histological sampling by crossection of blue mussels (Howard and Smith, 1983). However, some important organs are overlooked by these guidelines. We thus decided to add the adductor muscle in our screening. For histology, the samples were fixed in Davidson’s seawater fixative for 24-48 h before dehydration by series of ethanol baths to absolute ethanol concentration and subsequently embedding in paraffin blocks. The samples were sectioned for screening using routine staining with Hematoxylin Eosin (HE). The stained sections were investigated by light microscope, Leitz Weitzlar, Dialux 20EB, Germany as well as by phase microscopy, Nikon Labophot: Tokyo, Japan. Photos were taken using the same setup as described above.

Tissue fluid smears were collected on glass for cell morphology (cytological investigations) from digestive gland, hemolymph (withdrawn from adductor mussels by syringe) and also from biopsies of adductor muscles. Once dry, the smears were kept dark in boxes until usage.

### 2.5. CARD-FISH and confocal microscopy

To find DNA probes specific to *H. nelsoni*, we used a probe design pipeline available here: https://github.com/MiguelMSandin/oligoN-design. In brief, this pipeline requires a database of non-target sequences which we built using curated 18S rDNA sequences available in PR2 (https://pr2-database.org). This database was supplemented with a selection of marine bivalve diversity obtained from GenBank, representing potential marine bivalve hosts. This custom database is available in the Supplementary file: NonH_nelsoni_MarinMatrix2.fasta. We selected a probe specific to *H. nelsoni* which we refer to as Hex1: 5’-GCAGCTACCCTGTTATGA-3’. As negative control, the reverse complement of this sequence was used: Hex1-nose: 5’-TCATAACAGGGTAGCTGC-3’. We also used a general eukaryotic probe as positive control, EUK1209R 5’-GGGCATCACAGACCTG-3’ (Piwosz et al., 2021). All three probes were conjugated with Horseradish peroxidase (HRP) and synthetized by Biomers.net (Ulm, Germany).

To test the method for the identification of *Hexamita* spp., we first used our cultured trophozoites (M2-6738-22) and adapted previously published CARD-FISH protocols (Rodríguez□Martínez et al., 2022). First, we used 25mm diameter filters with pore sizes of either 0.2 µm or 5.0 µm (Track-Etch membranes, Nuclepore^TM^, Whatman^TM^, Cytiva, Buckinghamshire, UK). The cells were either fixed direct on the filters or transferred to PBS, and then prefixed before filtration in 4% marine formaldehyde (Davidsons marine fixative) for 1 hour. The filters were rinsed by washing in MilliQ water, air-dried and stored at -20°C until further processing. To decrease background signal, we also used cultured cells attached to glass slides. The culture was washed in PBS and mixed with eggwhite as described by Mazencova et al 2023. Three paralell drops of 5-10 µl was spread to form three sequential smears on the same glass, using small cover glasses (12×12 mm) for thin spread of fluid and instant drying. The smears were postfixed by Davidsons marine fixative in cold room (2-6 °C) for 8-12 hours before short washing in PBS, followed by MillQ-wash and air drying.

Probe hybridization was performed for 3 hours or overnight. To reduce background signal, two washing steps were extended to 30 minutes at 37°C, under slow shaking. The TSA step was prolonged to 1hour, incubating the samples for 1 hour. The fixed cells were kept frozen before analysis on an inverted fluorescence light microscope (Nikon Eclipse Ts2R) using a TIS Camera USB 3.0 Monochrome 1/1.2 CMOS and on an inverted confocal microscope (Leica DMi8) with Stellaris 5 scan head. For the latter the cells were analyzed using either: a) dry plan apochromatic 10x/0.40, b) dry plan apochromatic 20x/0.75, and for extra DIC functionality c) plan apochromatic 40x/1.25 glycerol emulsion and d) plan apochromatic 63x/1.40, oil emulsion.

In addition to cultured trophozoites, we also applied CARD-FISH to tissue sections. Histological sections were deparrafined and rehydrated in PBS during 1-2 hours before applying the CARD-FISH protocol described in (Blazejak et al., 2005) with the following modifications. In the prehybridisation step for paraffin-embedded samples, we increased the proteinease K concentration to 5 µg/ml and the incubation time to 10 minutes at 37°C. To decrease the background signal, we lowered the probe concentration (25 ng/µmol) and prolonged the washing steps to 2×30 minutes. We also increased the TSA incubation step to one hour at 46 °C in darkness using the amplification solution mixed with Alexa Fluor 488.

## 3. Results

### 3.1. Histopathology revealed cryptic lesions in adductor muscle and mantel of blue mussels

Wild blue mussels collected between 2017 and 2022 along the Swedish west coast were screened by histopathology and revealed undescribed lesions in the adductor muscle and mantel. In the adductor muscle, cross sections showed different grades of degenerative changes compared to healthy muscle anatomy (Figure 1A-B) where triangular or rectangular fibers normally occur. In the affected tissue, (Figure 1C-H) we observed mild, medium or widespread cellular swellings, loss of muscle fibers, vacuolization, cell-infiltration, and necrotic lesions. We graded the patterns of degenerative changes in the muscle fibers as mild cases (1C) when cell-infiltration led to low numbers of swollen fibers. We deemed cases as medium (1D) when observing 10 % to 50 % of unusual ballooned muscle fibers corresponding to cellular swelling. More severe pathologies were also observed (1E-F) with lytic destruction of the individual fibers, ultimately leading to vacuolization and fiber degeneration. In the more severe cases (1G-H), we observed a widespread degeneration of the cellular organization, with heavy vacuolization and oedemic infiltration, filling up seemingly empty areas between the muscle fibers with cellular debris or transparent fluid. In these areas with extensive destruction of muscle bundles, the sectioned organ was partially or totally lacking normal muscle tissues.

**Figure 1.**
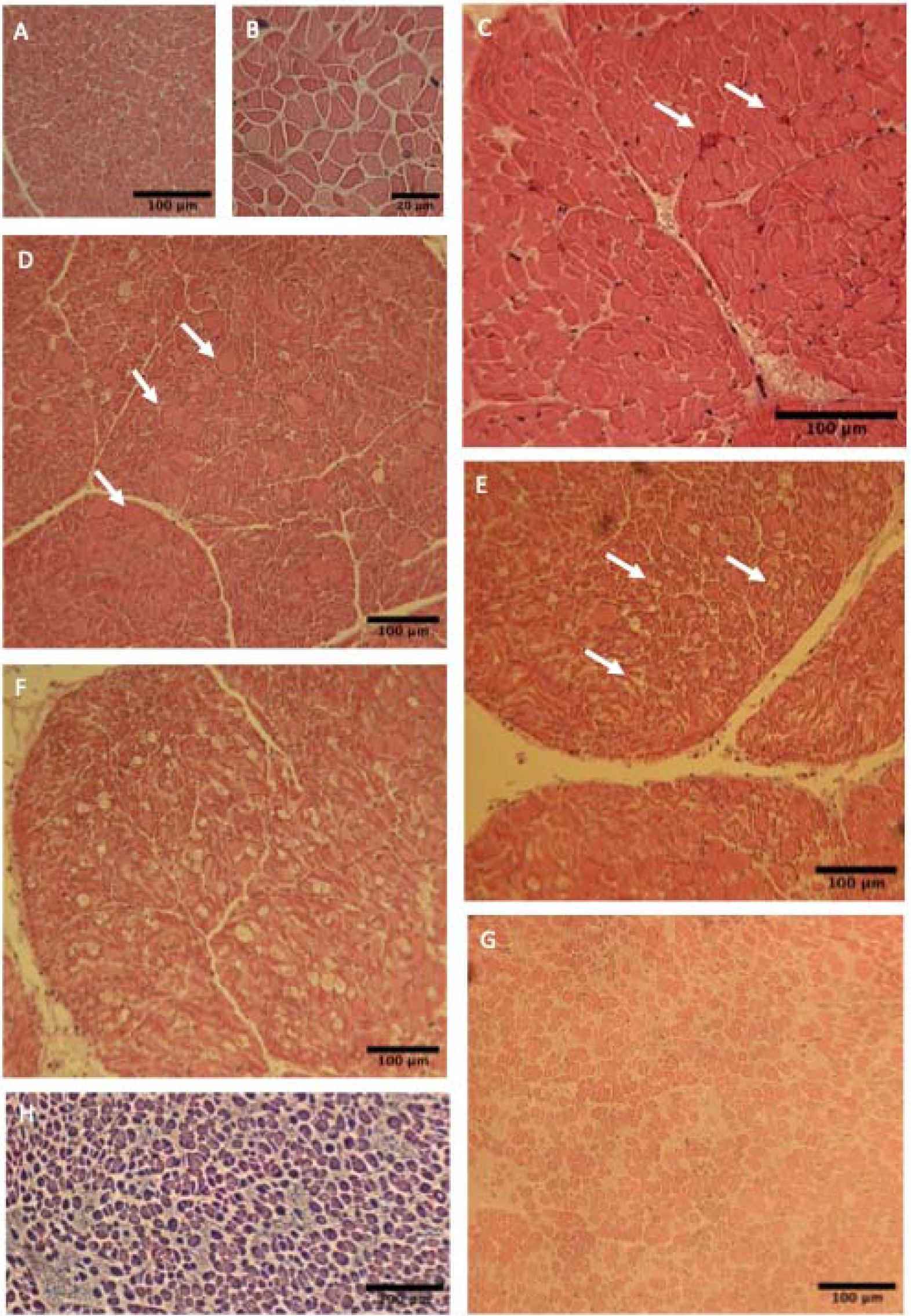
Histology adductor muscle *M. edulis*, blue mussel revaling degenerative changes and lesions in blue mussel host. A-B) Healthy i.e. normal triangular or rectangular cross-sectioned adductor muscle fibers. C) mild cellinfiltration and a few cases of ballooning fibers. D) multiple round fibers indicating increasing grade of cellular swelling. E) widespread pattern of vacuolization and muscle degeneration, mixed with ballooning fibers. F) Destruction of the individual fibers, increasing vacuoles and patterns of necrotized fibers. G) More than 50 % of the fibers transparent, empty or replaced with oedemic fluid. H) Phase contrast of the same area, clearly demonstrating the loss of normal muscle tissue.

In addition to the adductor muscle, the histological examination, also revealed degenerative changes in the mantel and specifically the gonad follicles as well as nutrient depot, ie. adipose glands (ADG clusters). In the case of female ovary follicles, the degenerative changes was sometimes also affecting the oocytes, where these seemed to collapse, loosing their normal egg shape (Supplementary Figure 1).

### 3.2. Enrichment of flagellates from blue mussel muscle biopsies and light microscopy

We hypothesized that protozoan parasites not monitored by routine surveillance programs were causing the degenerative lesions observed in the adductor muscle and mantel. To test this hypothesis, we performed additional small biopsies from the adductor muscle on samples collected in November 2022. From these biopsies, we were able to isolate flagellated cells from two mussels collected from the Grebbestad municipality (coordinates:11.13,4650 x 58.42,6299), after incubation at low temperature (4^°^C) in Marine EMEM cell culture medium for two months. One isolate (M2-6738-22; deposited*:pending*) has been maintained as a continuous culture since isolation. A few similar incubation experiments were performed using biopsies from material collected in previous years (2018/19), from mantel, adductor muscle (Table 1). However, while we observed the proliferation of similar flagellates, the enrichments were not stable and often collapsed suddenly or were overgrown by fungal or bacterial contaminants.

**Table 1.**
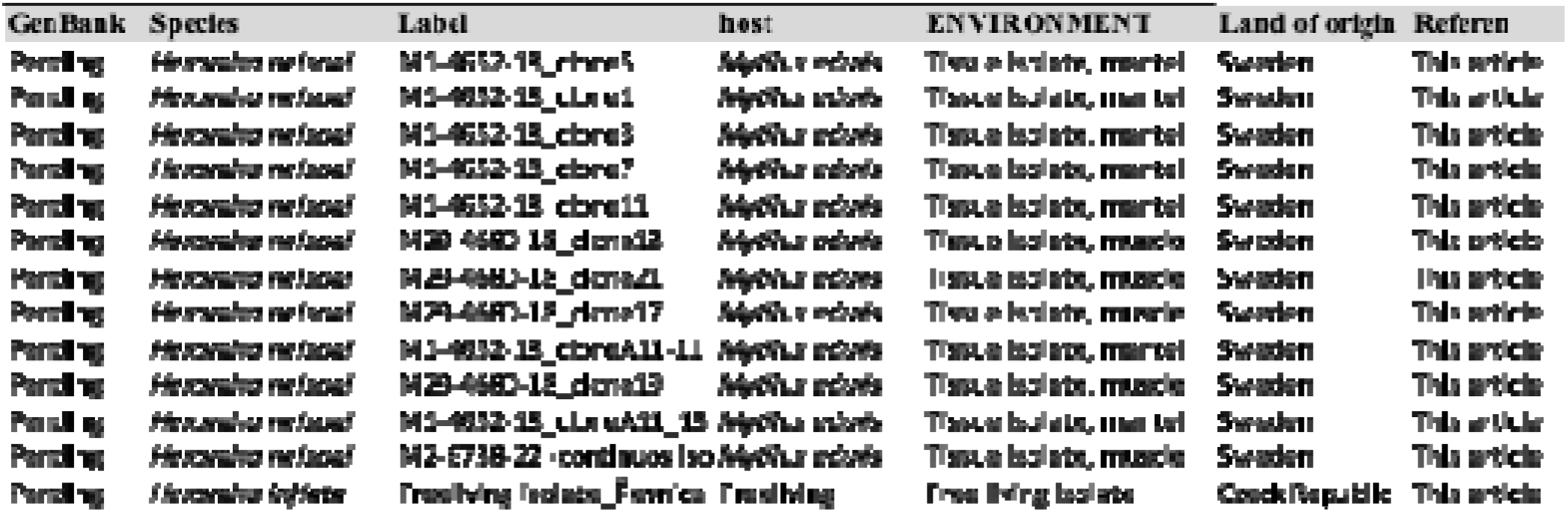
Isolates of Hexamita spp. used in this study.

Upon closer inspection of isolate M2-6738-22 by light microscopy, we observed broadly similar cell characteristics as to the original description of *Hexamita nelsoni* trophozoites, isolated from the American oyster *Crassostrea virginica* (Schlicht and Macking, 1968). The cells harbored 2 nuclei placed anteriorly, 6 anterior flagella (AF), and 2 long recurrent posterior flagella (PF) close to the cell surface positioned within lateral axostyles or cytostomal tubular channels (Figure 2, Supplementary Figure 2). At approximately 2/3 of the body length, these opened up into a narrow cytostomal-like groove (Figure 2A-D). These narrow groves reminded us of “old style la crosse ballgame bats” used to catch and throw spherical balls. We observed the trophozoites preying on bacteria via movements of the anterior flagella (Figure 2B-D, Supplementary movie 1). The bacteria were collected within the grooves, and then transported forward and enclosed into vesicles that we interpret as food vacuoles (Figure 2B-C, F-G). In the supplementary movie, these food particles can be seen in a chamber connected to the cytostome directly beneath the nuclei.

**Figure 2.**
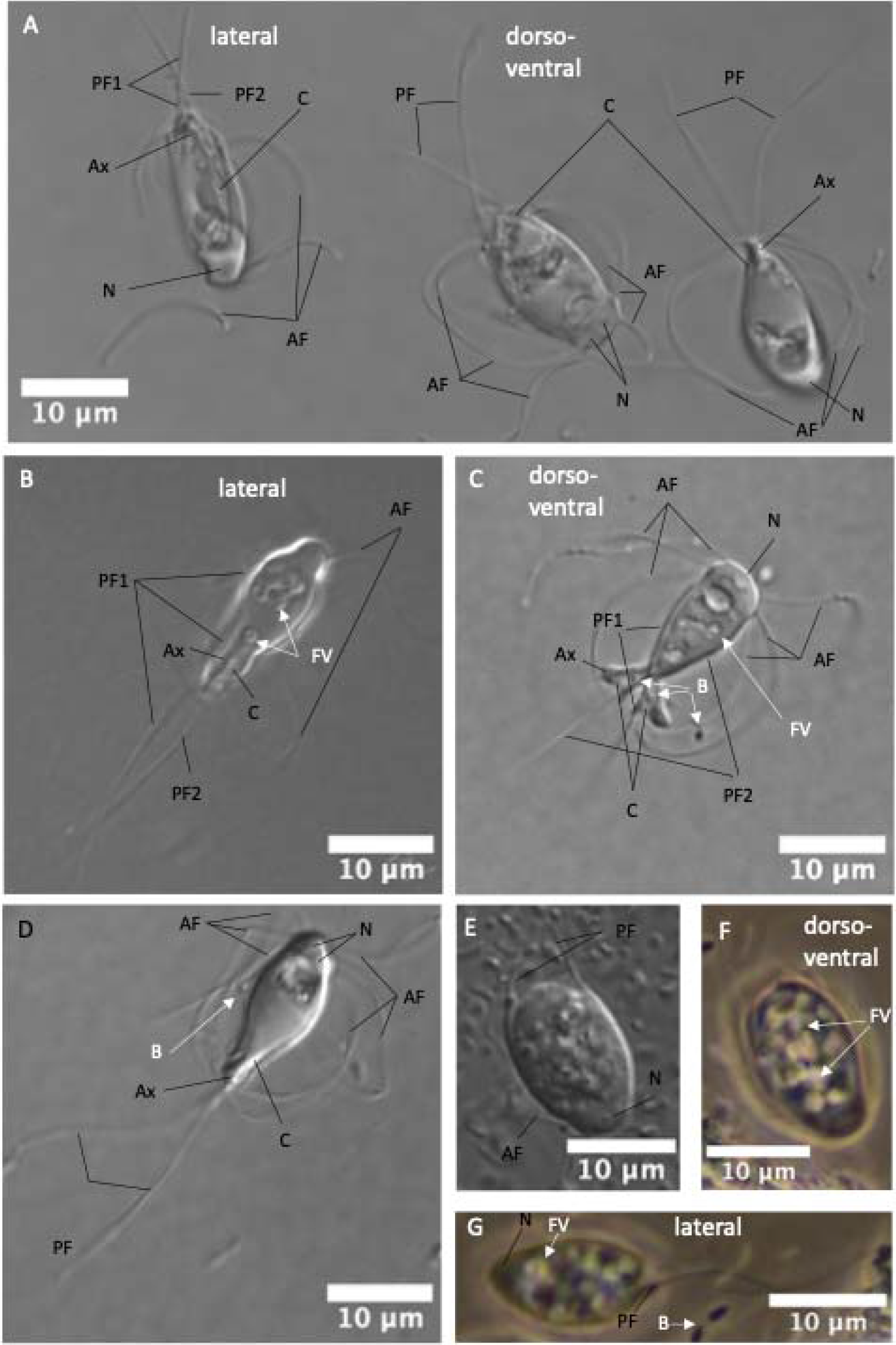
Cytology, trophozoites in cell culture observed by: differential interference contrast (DIC) 2A-E, and phase contrast (PH) 2F-G. Note the flexible lateral pairs of anterior flagella, in total 6, and the two straight, recurrent and posterial tracing flagella (PF), Figure 2A-D. show dorsoventral fattened cells, filled by food vacuoles, observed by DIC (2E) and PH (2F) and in lateral view (2G).

**Figure 3.**
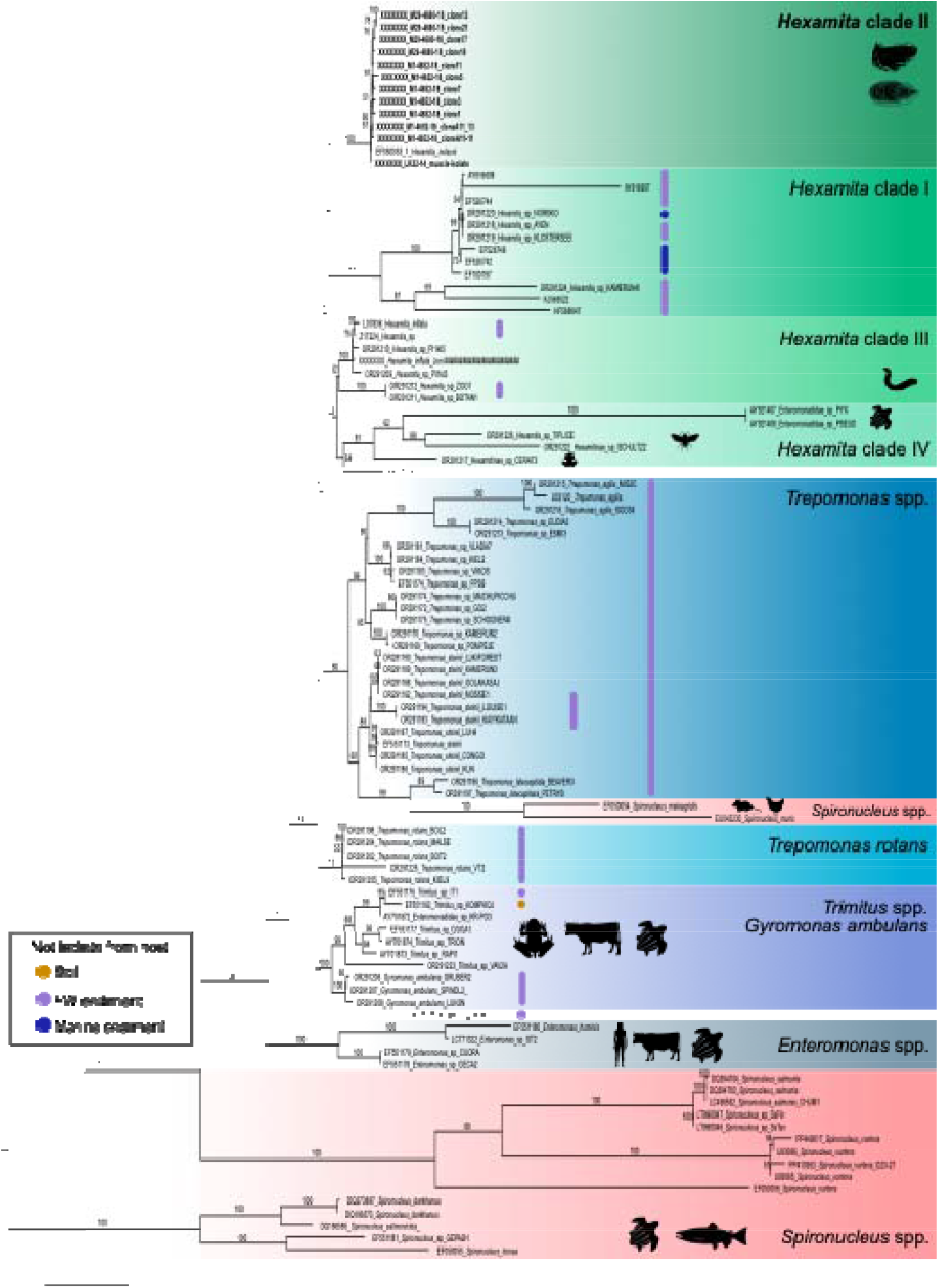
Maximum likelihood phylogenetic tree based on the 18S rDNA gene showing the placement of the blue mussel-isolated sequences corresponding to *H. nelsoni*, in a broaded context of diplomonad sequences. The number on branches is bootstrap support based on 100 replicates (bootstrap support below 50 the taxon name are listed. Known hosts are shown using pictograms. When not host-associated, colored dots are used to indicate the source environment. The tree was rooted on Spironucleus sequences. The scale bar represents the estimated number of substitutions/sites.

Live cells measured around 15.0 (+/-1.23) µm in length and 9.4 (+/- 1.55) µm in width. Cells fixed in Davidsson marine fixative and mounted with egg white were generally longer and slender (Table 2). The trophozoites were swimming rapidly with sudden directional shifts and with a continuous spinning of the body along the longitudinal axis (Supplementary movie 2). The cell shape was variable, not always taking the typical pyriformic form of *H. nelsoni* as described in Schlicht and Mackin (1968). In swimming cells, we also observed forms ranging from ellipsoid, oblong-ovoid, or ovoid, with a truncated, tridental or concave posterior ending (Figure 2, Supplementary Figure 2). A small number of spherical cyst-like cells were also observed enclosing inner double or quadruple nuclei (Supplementary Figure 2J-L). In our cell culture, we also observed multi-flagellated ballooned trophozoites appearing to enclose two or more cells moving in every direction. These enlarged cells, measuring up to 40µm in length when moving in opposite direction, indicated binary fission and cell division enclosing inner trophozoites (Supplementary Figure 3).

**Table 2:** measurements of live and fixed cells on glass (Davidsons marine fixative)

| **Table 2** | **Length** | **stdev.** | **Width** | **stdev** | **L/W** | **stdev.** | **Nr** |
| --- | --- | --- | --- | --- | --- | --- | --- |
| Live trophozoite | 15 | 1,2 | 9,4 | 1,6 | 1,6 | 0,2 | 38 |
| Egg white/Fixed | 18,2 | 2,1 | 7,4 | 1,4 | 2,4 | 0,4 | 20 |

### 3.3. Phylogenetic position of trophozoites isolated from blue mussels

To support the hypothesis that the flagellated cells isolated from blue mussels are related to Hexamitinae, we sequenced the 18S rDNA gene from the new isolate M2-6738-22. We also sequenced 11 clones from two previous isolates (Table 2) which were all nearly identical (sequence similarity = XX% *pending*). All clones were placed in a phylogenetic context using a Maximum Likelihood (ML) tree reconstruction method including a broad diversity of related diplomonad sequences (Figure 3). The clones clustered closely to the endobiotic *H. nelsoni* from oysters (accession number EF050053), and the new blue mussel isolate M2-6738-22, in a maximally supported clade (bootstrap = 100%) named “Hexamita clade II” for consistency with Mazancová et al. (2023). As expected from previous work (Jørgensen et al 2007, Mazencova et al 2023), the genus *Hexamita* was recovered polyphyletic, split into 4 groups referred to as clades I to IV and including *H. inflata* and other free-living isolates as well as host-associated isolates (Figure 3). *Hexamita* clade I received strong support (bootstrap = 100%), as was the case–albeit slightly lower support–for the grouping of *Hexamita* clade III and IV (bootstrap = 92%). However, both clades III and IV individually received only weak support. Our tree also included the free-living genus *Trepomonas* (also recovered polyphyletic), as well as the endobiotic or free-living genera *Trimitus* and *Gyromonas* and 3 paraphyletic parasitic clades of *Spironucleus*. In general, the relationships among the main groups in the tree received only moderate to low support.

### 3.4. Connecting parasite and host with CARD-FISH

We used CARD-FISH to identify a possible link between the *H. nelsoni* isolate from blue mussels and the lesions observed by histology in the adductor muscle and mantel. A specific probe (Hex1) was designed to target the rRNA in the *H. nelsoni* clade and first tested on the M2-6738-22 culture. The probe localized to cells attached and fixed on two different filter sizes (5µm and 0.2µm). Many of the cells appeared to be in different stages of division, as seen by the high number of large, multinucleated cells (Figure 4A-J). The staining pattern is more dense around the nuclei and in an extensive network throughout the cells, probably due to hybridization of ribosomes at the endoplasmic reticulum (Figure 4A-D).

**Figure 4:**
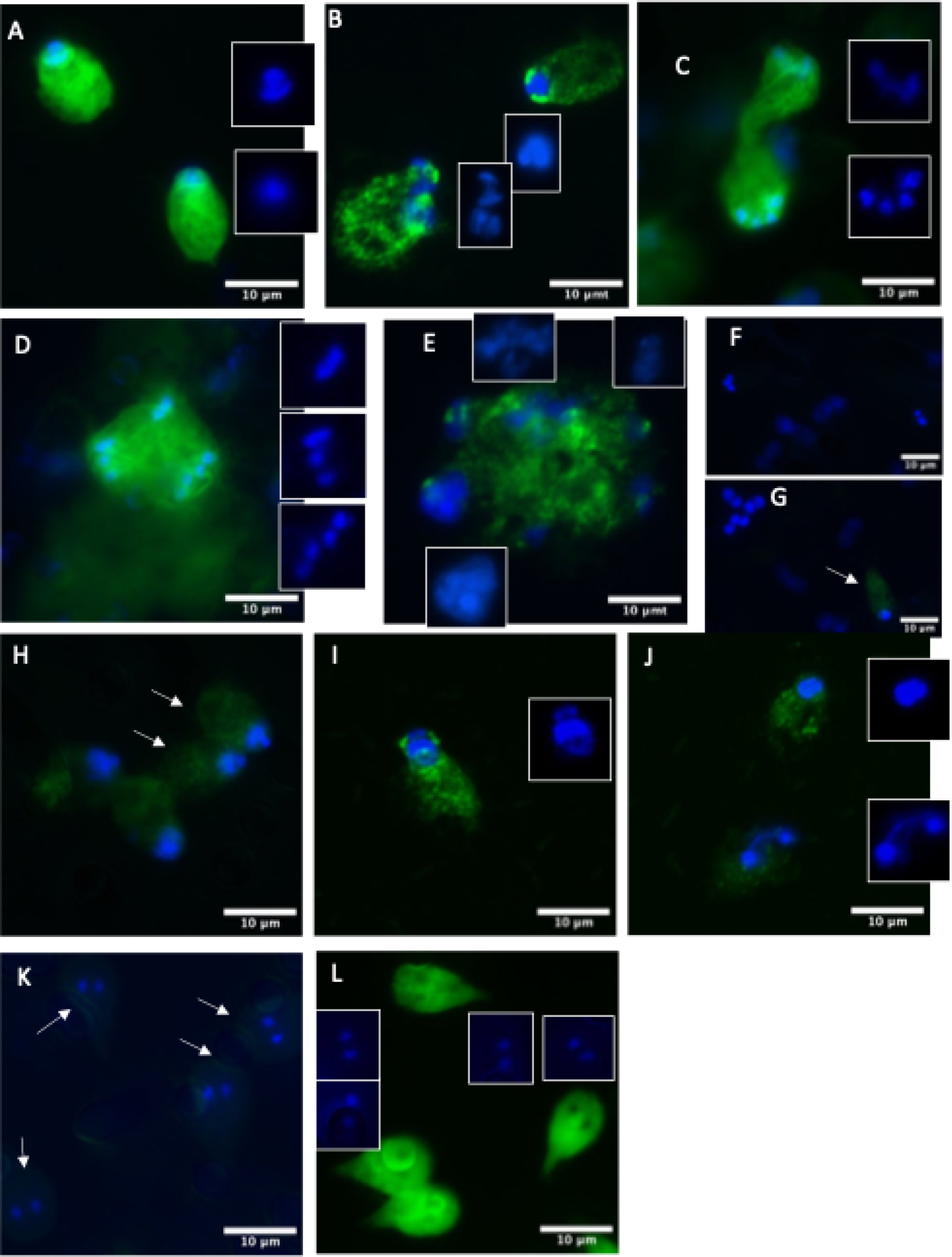
CARD-FISH staining of cell culture (on 5µm filters) of *Hexamita nelsoni* isolated from *Mytilus edulis*, LK22-14, using 50ng/µl probe concentration. Figure A-E: show Hex1 probe signal, also indicating the DAPI signal (enclosed in white squares). Figure F-H controle: absence of signal, indicate cells stained without adding Hex1 probe (F: No probe) and G weak signal (G-H: NS probe 50ng). Evaluate probe at lower concentration I-J, Figure I: 25ng signal, J weak signal. Figure k-L Diplomonad control: Giardia very weak signal Nsprobe, compared with reference probe: genEUK Figure L

To test for probe specificity, we first used a negative control consisting of the hybridization buffer without the probe, which did not provide signal (Figure 4K-L, Supplementary Figure 4A-C). We also used an alternative negative control consisting of the reverse complement sequence of Hex1. When used at the same initial concentration as Hex1 (50ng/µl), we noticed a weak background signal (Supplementary Figure 4D, F). To reduce this background signal of the reverse complement probe while maintaining specificity of Hex1, we tested two dilutions of the probe (25ng/µl and 10ng/µl). This showed that at 25ng/µl, the unspecific background signal was reduced to a minimum while Hex1 continued to show the expected specificity (Supplementary Figure I-L). Finally, we used an additional negative control by testing the Hex1 probe on a related but different diplomonad culture (*Giardia intestinalis* isolate WB, assemblage A), and contrasted this with the signal obtained with a general eukaryotic probe (GenEuk). Although we detected a very weak signal with Hex1 (Supplementary Figure 4M-N), the GenEuk probe resulted in a much stronger and clearer signal (Supplementary Figure 4O), again around the nuclei and in an ER-like pattern. Overall, our testing of the probes and hybridizing conditions indicate that Hex1 provides adequate specificity.

We then applied the Hex1 probe to sections of adductor muscle and mantel biopsies of blue mussels. In a mild case infiltration of the adductor muscle, we observed multiple positive elongated cells positioned around healthy muscle fibers (Figure 5A), with nuclei occurring in extreme anterior position (Figure 5B), similar to our observations of the cell culture (Supplementary Figure 1F and 2H). Corresponding observations of *Hexamita*-like cells positioned around muscle fibers were also made by light microscopy based on traditional Hematoxylin-Eosin (HE) staining (Figure 5C). In some cases, multiple Hex1 positive cells surrounded the muscle fibers (Figure 5D), or sometimes expanded to enclose single fibers (Figure 5E), an arrangement also revealed by histology (Figure 5F). We also observed larger cells with spherical or rounded shape embedded within the fibers (Figure 5G-H), which resembled cells also observed by HE staining (Figure 5I). In Figures 5J and K a double nucleated cell characteristic for Hexamitinae is seen. Figure 5L show large cells with nuclei in outreach position infiltrating around the adductor fibers.

**Figure 5:**
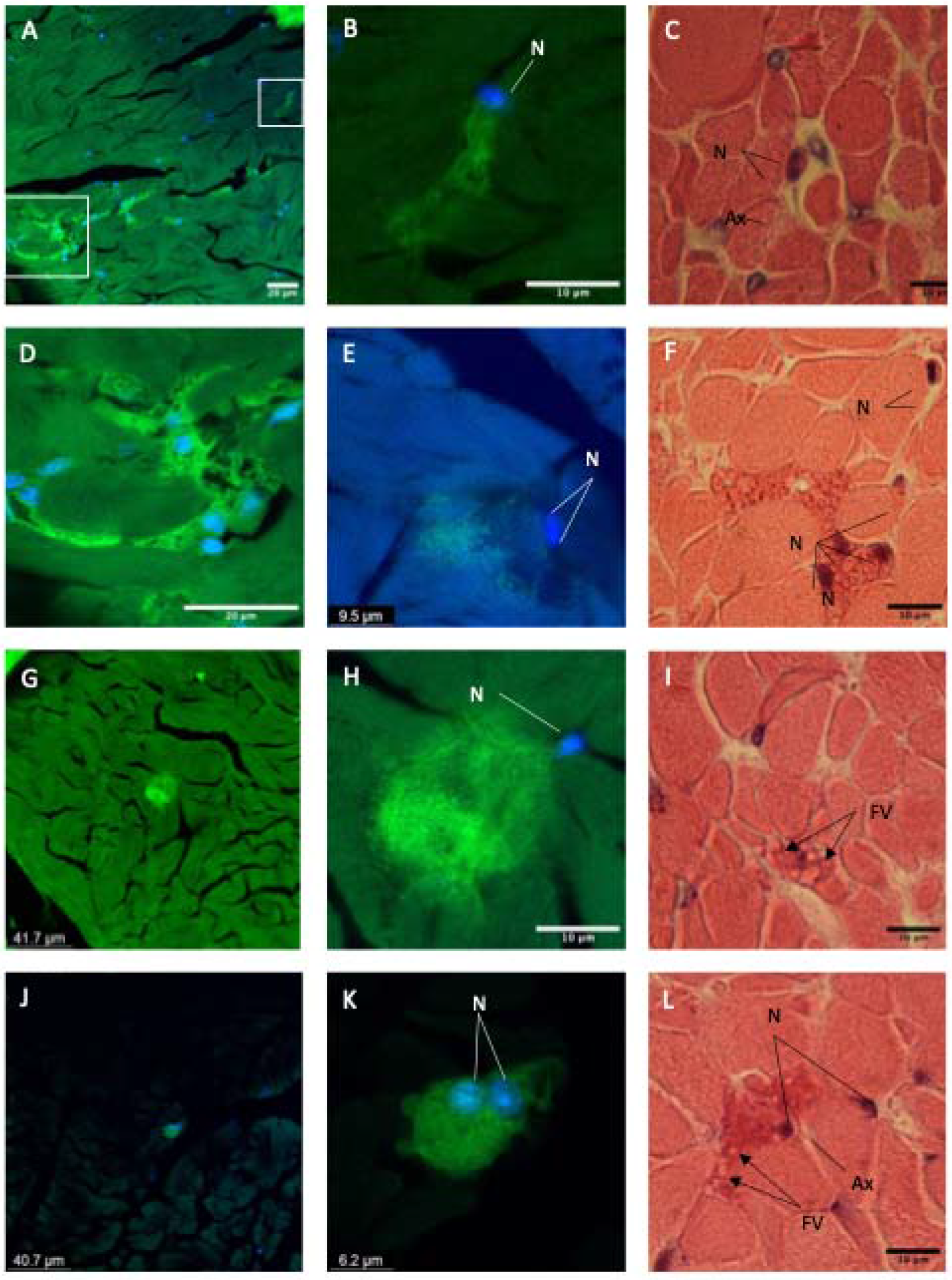
CardFISH hybridisation on *Mytilus edulis*, bluemussel, abductor muscle (A-B, D-E, G-H, J-K) and Histology, HE stained muscle biopsies (C, F, I, L) showing signs of *Hexamita* like cells observed within and between abductor muscle fibers. In Figure 5E, F, J and K, double nucleated cells, characteristic for Hexamitinae is seen. Figure 5B, E, F, H and L show cells with single/double nuclie in anterior or outreach position. Figure C and and an axostyle like structure infiltrating around the adductor fibers. Abrreviations N=nuclei, FV = food vacuole, AX = axostyle like structure

In the mantel, we detected individual Hex1 positive cells within the adipose glandular tissue (ADG) (Figure 6A-B). The ADGs, responsible for storage of absorbed nutrients in fattening mussels, are seen as pink to red clusters of aggregating cells between gonad follicles (Figure 6C). In some of these clusters, an infiltration of multiple small Hex1 positive cells were observed (Figure 6D-E). Corresponding patterns of degenerative changes withing the ADG cells were also detected with HE staining of mantel tissue (Figure 6F). Some of the ADG clusters, situated close to, or directly outside of the ovary follicles, were entirely positive to the Hex1 probe. Within these clusters, multiple double nuclei were observed, supporting our inference that these are *H. nelsoni* (Figure 6G-H). Similar large eosinophilic cells with inner segmentation are indicated in figure 6I-J. Within the ovary follicles filled with oocytes we also observed infiltration of Hexamita like cells by Card-FISH (Figure 6K-L) also observed by normal histology (Figure 6M).

**Figure 6.**
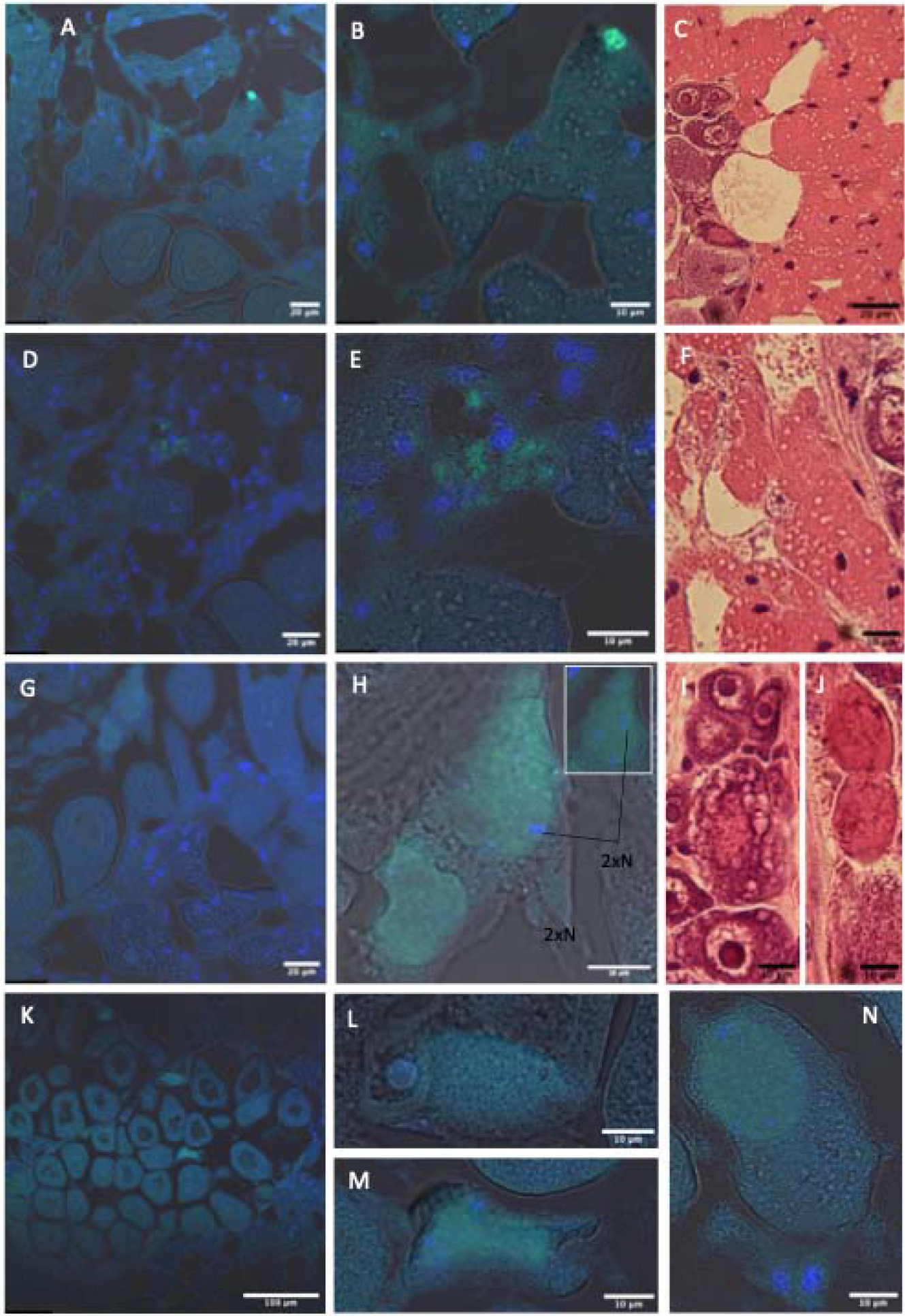
Catalyzed Reporter Deposition Fluoroscence in Situ Hybridization (CARD-FISH) of mantel and gonads. A-B) Adipose glandular cells with single positive cells. C) In HE stained mantel these ADG cells are in close contact with gonad follicles. D-E) Degenerative changes in ADG cluster with positive stained cells. F) Pattern of vesicular degenerative changes. G-H) Cellcluster of positive cells, seen in outer part of gonad follicle. Note double nuclei (DAPI blue) observed within these. I-J) Corresponding gonad aree where clear vacuolisation can be observed within ovary oocytes. In the J small enclosed spherical vacuoles or µ-cells are seen. K-M) Infiltration of gonad of hexamita positive, cells resembling collapsed oocytes enlosing inner Hexamita cells. N) Oocyte with Hex1 positive sphere indicating also four Dapi positive nuclei, closely together.

## 4. Discussion

This study reports the first discovery of the diplomonad flagellate *Hexamita nelsoni* in blue mussels (*Mytilus edulis*). Investigating cryptic lesions in specimen collected as part of the routine national surveillance program of bivalves in Sweden targeting *Marteilia* spp., our combined molecular and microscopic observations (both light and targeted confocal microscopy) show that *H. nelsoni* is associated with pathological tissue in the gastro-intestinal system and gonads. Using a newly established culture of the blue mussel strain, we propose a morphological description that confirms some unique morphological characteristics only reported in the original species description from oysters (Schlicht and Mackin 1968).

The first species description in the Hexamita genus, corresponding to *H. nelsoni*, was based on trophozoites isolated from American oysters *(Crassostrea virginica)* (Schlicht & Mackin 1968). These flagellated cells bore 8 flagella–6 anterior and 2 posterior–and measured about 8 x 15 µm. The posterior ending of these trophozoites was shown as rounded, concave or slightly posteriorly projecting, i.e. “flaring”. In most cases, two nuclei could be discerned but often appeared as one large dark staining region in a very forward position (Schlicht & Mackin 1968). This morphological description was similar to the single figure published earlier by both Scheltema 1962 and Mackin et al. 1951, but size differences were noted. Our blue mussel isolate, the first from this bivalve group, is very similar to the oyster-isolated type species in its general morphology and in size. The overall morphological similarity was corroborated by nearly identical sequences to the only *H. nelsoni* 18S rDNA sequence available, derived from cells isolated from Norwegian flat oysters (Jørgensen and Sterud 2007). Our observations based on phase contrast and DIC also confirmed a unique characteristic of *H. nelsoni* compared to other Hexamita isolates–all free living–which was already proposed in the original species description. Indeed, we interpret the posterior ending with two flaring parts in starved and slender trophozoites (Figure 1C, Supplementary Figure 1A-C) as homologous to the flared or bell-shaped axostyle posterior ending in Schlicht and Macking (1968).

The observation of flagellates related to *H. nelsoni* have a long, although sparse, history in the literature. While several free-living isolates of diplomonads have recently been described (Mazancova et al 2022), including the free-living species *H. inflata* (Lloyd and Williams, 2014), it is the host-associated lineages that have concentrated most of the attention. The first report of Hexamita-like cells, isolated from European flat oysters (*Ostrea edulis*), date back to 1882 (Certes, 1882), nearly 80 years before a formal species description became available (Schlicht & Mackin 1968). Since then, similar cells have been observed to infect various animals, including oysters, fishes, amphibia (Woo & Poynton 1995), birds and mice (Hinshaw et al 1938, McNeil et al. 1941 and Meshoker 1969), but molecular data are often lacking to confirm the exact identity of the cells. In bivalves, the presence of corresponding cells has been noted in other oyster species such as the Olympia oysters (*O. lurida*) (Stein et al 1961) and the American oysters (*Crassostrea virginica*) (Scheltema 1962). In the majority of cases of other Hexamitinae, infecting aquatic vertebrates, the cells have been only observed in the gastro-intestinal system, where they can cause enteritis and malnutrition leading to lethargy, starvation and eventually mortalities. (Fard et al., 2007; Jørgensen and Sterud, 2007b; Levy et al., 2015; Sterud et al., 1997; Williams et al., 2011). In a cohabitation trial experiment, Olympia oysters indicated–although inconclusively–higher mortality when exposed to *Hexamita* compared to the unexposed control population (Stein et al 1961). The American oysters showed a seasonal variation in the prevalence of *Hexamita* in Delaware bay (New Jersey, USA), with a maximum infection reaching up to 90% prevalence in early spring when the bay experienced the lowest water temperatures with detrimental effects on the phenotype leading to increased mortality (Scheltema 1962, Feng and Stauber 1968). In general, however, the effects on the hosts of these *Hexamita*-like cells are poorly understood and often inconclusive, and it has remained unclear whether this organism is parasitic or commensal and what are the routes of infection.

Interestingly, our observations of *H. nelsoni* in different organs, not just the gastro-intestinal system where it has been most often reported before from various hosts (Mazencova 2023, Woo, Poynton 1995, Fard et al 2007, Gallani et al 2016), indicate that in blue mussel at least a more systemic infection is possible. Combining classical histology with targeted CARD-FISH for *H. nelsoni*, we found positive cells in the adductor muscle, wrapping around individual or multiple fibers, within ADG clusters in the mantel (responsible for nutrient storage), and in the gonad follicles (Figure 6). The possibility of a systemic infection of *Hexamita* cells in other bivalve hosts is not unprecedented, but these early studies in oysters lacked molecular identification. Still, flat oysters in the Netherland were reportedly systemically infected by flagellated trophozoites with 3 pairs of anterior flagella and two posterior trailing flagella (Mackin et al. 1951), which corresponded to the later species description of *H. nelsoni* (Schlicht and Mackin 1968). Other organs (digestive glands, hemolymph within blood vessels, mantel, ovaries and adductor muscle) in the European flat oysters were also reportedly invaded by cells lacking flagella, sometimes intracellularly, but which were interpreted as different stages of the same protozoan infection (Macking et al 1951). From these observations, several conclusions were drawn, which are congruent with our more partial findings in blue mussels of *H. nelsoni* in the adductor muscle, mantel, and gonads: 1) *H. nelsoni* in bivalves are parasitic protozoans found mostly within blood vessels and not in connective tissue, gathering in sinuses or basal membranes of stomach and intestine. 2) Trophozoites can aggregate in abscesses, sometimes only within necrotic tissue. 3) The infection can spread secondarily to gonads with degeneration and liquefied tissues or inclusion bodies within ovaries. 4) The parasitic infection leads to degenerative changes in both stomach and digestive gland, with cells accumulating within the epithelium (Mackin et al 1951).

While it remains to be seen how *H. nelsoni* impacts blue mussels health and recruitment, and whether it is an opportunistic parasite that becomes a risk only in unfavorable conditions, it is worth noting that related parasites of the genus *Spironucleus* have been better documented to systemically infect their hosts. This is for example the case for *S. vortens*, which is described from the liver, spleen, and kidney of fishes (Paull and Matthews, 2001; Sangmaneedet and Smith, n.d.) confirming the systemic nature of this pathogen (Paull and Matthews, 2001). *S. salmonicida*, infecting salmonids, is another example that is specifically recognized as a systemic pathogenic parasite causing sudden and severe mortalities during winter seasons (Kent et al 1992, Williams et al 2011, Sterud et al 2003, Poppe et al 1992, Guo and Woo 2004, Sterud and Jørgensen 2006).

The finding of *H. nelsoni* infecting blue mussel and causing lesions not previously described is significant because these organisms are cornerstone to the shellfish industry. Although heavily monitored by governmental agencies for a range of listed pathogens, blue mussels and other bivalves potentially host other untested parasites that can, under specific circumstances, increase mortality in farmed but also natural populations. It is thus important that more efforts are taken to look at reoccurring cryptic lesions without obvious pathology in bivalve hosts, in order to get a more comprehensive picture of the complex causes of decreased fitness in economically important species. Here, we have showed that *H. nelsoni* is responsible for such cryptic lesions in a new host, but it is most likely that continuing searching for cryptic parasites will reveal a large diversity of parasites.

## Author contribution

AA and FB designed this study. AA performed health inspection and sampling. AA and JJH performed cell culture and lab work. AA, FB, JJH analyzed data and prepared figures. AA, JJH, SS and FB wrote and edited the paper.

## Acknowledgment

We thank the Swedish Veterinary Institute SVA and the department of Fish for support with material and cell media as well as the histopathology lab for technical support in implementing the CARD-FISH method. We are grateful to members of Burki lab (Yash Pardasani, Nina Poll and Anne Walraven) for help with probe design, phylogenetic analyses, and applying the CARD-FISH methodology to cell culture and histology. We also want to thank the Swedish Agency for Marine and Water Management for supporting the surveillance of wild bivalve, and the oyster fisheries and aquaculture facilities enabling the collection of blue mussels.

## Financial support

This work was supported by SciLifeLab, a Formas Future Research Leader grant (2017-01197), and a Swedish Research Council VR research project grant (2017-04563) all available to FB.

## Notes

### Competing Interest Statement

The authors have declared no competing interest.

