## Supplementary Table 1 for "First report of *Hexamita nelsoni* in blue mussel (*Mytilus edulis*): morphology, phylogeny, and host-parasite interaction"

**Table 2.** Genbank, reference sequences, 18s rDNA gene partial

| GenBank numb | Species | host | ENVIRONMENT | Land of origin | Referen |
|---|---|---|---|---|---|
| PP449017 | Spironucleus vortens | *Symphysodon discus* | Digestive tract of discus fish | Poland | Pastuszka,A 2024 |
| PP410363 | Spironucleus vortens | *Herichthys cyanoguttatus* | Blue Texas Cichlid | USA | Cagle L.A. 2024 |
| U93085 | Spironucleus vortens | ? | ? | Canada | Keeling and Doolittle 1997 |
| U93086 | Spironucleus vortens | ? | ? | Canada | Keeling and Doolittle 1997 |
| EF050056 | Spironucleus vortens | *Leuciscus idus* | Digestive tract of ide | Norway, Oslo river | Jorgensen,A. and Sterud,E.2007 |
| DQ394704 | Spironucleus salmonis | *Oncorhynchus mykiss* | Digestive tract of rainbow trout | Germany | Fard 2006 |
| DQ394703 | Spironucleus salmonis | *Oncorhynchus mykiss* | Digestive tract of rainbow trout | Germany | Fard 2007 |
| LT990046 | Spironucleus sp. SsTsn | *Perccottus glenii* | Intestine of Amur sleeper fish, Egorievsk fish farm | Russia | Denikina,N.N.2018 |
| LT990047 | Spironucleus sp. SsTsn | *Perccottus glenii* | Intestine of Amur sleeper fish, Egorievsk fish farm | Russia | Denikina,N.N.2018 |
| LC496562 | Spironucleus salmonis CHUM1 | *Oncorhynchus keta* | Intestine of Chum salmon | Japan | Mizuno,S. and Urawa,S. 2020 |
| EF050055 | Spironucleus torosa | *Gadus morhua* | intestine of Atlantic cod | Norway? | Jorgensen,A. and Sterud,E.2007 |
| DQ186597 | Spironucleus salmonicida | *Salvelinus alpinus* | blood from farmed arctid char | Norway, fiskfjord | Jorgensen,A. and Sterud,E.2006 |
| DQ186585 | Spironucleus salmonicida isolate 2 | *Salmo salar* | inner organ of farmed atlantic salmon | Norway, Alta | Jorgensen,A. and Sterud,E.2006 |
| DQ186573 | Spironucleus barkhanus isolate 1 | *Thymallus thymallus* | Digestive system of grayling | Norway, Glomma | Jorgensen,A. and Sterud,E.2006 |
| DQ273887 | Spironucleus barkhanus | *Thymallus arcticus* | Digestive system of grayling | Canada: Enadai Lake, Kazan Ri | Jorgensen,A. and Sterud,E.2006 |
| EU043230 | Spironucleus muris | Mammalian | Intestina tract? | Canada? | Kolisko M. et al 2008 |
| EF050054 | Spironucleus meleagridis | *Meleagris gallopavo* | intestinal tract turkey | Norway? | Jorgensen,A. and Sterud,E. 2007 |
| EF050053 | Hexamita nelsoni | *Ostrea edulis* | Intestine of flat oyster | Norway | Jorgensen,A. and Sterud,E. 2007 |
| L07836 | Hexamita inflata | | Environment free living | | Leipe, D.D. et al.1993 |
| Z17224 | Hexamita sp. | | Environment free living | | van Keulen, H. et al 1993 |
| EF551178 | Enteromonas sp. GECA2 | *Geochelone carbonaria* | Fecal isolate red footed turtoise | Czech Republic?/Southamerica | Kolisko M. et al 2008 |
| EF551179 | Enteromonas sp. CUORA1 | *Cuora amboinensis* | Fecal isolate amboica box turtle | Czech Republic?/Southeast asia | Kolisko M. et al 2008 |
| EF551180 | Enteromonas hominis ENTEROII | *Homo sapiens* | Fecal isolate human | Czech Republic? | Kolisko M. et al 2008 |
| AY921408 | Enteromonas sp. PSEUD | *Trachemis scripta elegans* | Fecal isolate red-eared slider | Czech Republic?/Northamerica | Kolisko M. et al 2008 |
| AY921407 | Enteromonadidae sp. PYX | *Pyxidea mouhotii* | Fecal isolate keeled box turtle | Czech Republic? | Cepicka, I. et al. 2005 |
| AY701872 | Enteromonadidae sp. KR-PO3 | *Bos taurus* | Fecal isolate cattle | Czech Republic? | Kolisko M. et al 2005 |
| EF551182 | Trimitus sp. KOMPOJ | Environment, soil | Compost, Kojčice, soil | Czech Republic | Kolisko M. et al 2008 |
| AY701873 | Trimitus sp. RAPI1 | *Rana pipiens* | Fecal isolate from northern leopard frog | Czech Republic? | Kolisko, M. et. Al. 2005 |
| AY701874.1 | Trimitus sp. TRION | *Aspideretes gangeticus* | Fecal isolate from indian softshell turtle | Czech Republic? | Kolisko, M. et. Al. 2005 |
| EF551176 | Trimitus sp. IT1 | Environment, freshwater | Pond in Italy | Czech Republic | Kolisko M. et al 2008 |
| EF551177 | Trimitus sp. DOGA1 | *Dogania* sp. | Fecal isolate from softshell turtle | Czech Republic? | Kolisko M. et al 2008 |
| EF551181 | Spironucleus GEPA2H | *Geochelone pardalis* | Fecal isolate leopard turtle | Czech Republic?/Southafrica | Kolisko M. et al 2008 |
| EF551173 | Trepomonas steini LUH3 | Environment, freshwater | Flood, Vltava river, South Bohemia, | Czech Republic | Kolisko M. et al 2008 |
| EF551174 | Trepomonas sp.2 PPS6 | Environment, freshwater | Point Pleasant Park pond, Halifax, NS | Canada | Kolisko M. et al 2008 |
| EF551175 | Uncultured eukaryote CHES12 | *Chelodina* sp. | Fecal isolate snake necked turtle | Czech Republic?/Australia? | Kolisko M. et al 2008 |
| LC771322 | Enteromonas sp. ST2 | *Bubalus bubalis* | Fecal sample buffalo_Wainyapu_B3_2014 | Indonesia:Sumba, Wainyapu | Lacante,S.A et al. 2023 |
| OR291184 | Trepomonas sp.2 KIEL2 | Environment, freshwater | Freshwater sediment | Behrensdorf, DEU | Mazencova et al., 2023 |
| OR291183 | Trepomonas sp.2 VIKOS | Environment, freshwater | Freshwater sediment | Vikos, GRC | Mazencova et al., 2023 |
| OR291181 | Trepomonas sp.2 VLADA7 | Environment, freshwater | Freshwater sediment | DEU | Mazencova et al., 2023 |
| OR291172 | Trepomonas sp.3 GO2 | Environment, freshwater | Freshwater sediment | Crete, GRC | Mazencova et al., 2023 |
| OR291174 | Trepomonas sp.3 MACHUPICCHU | Environment, freshwater | Freshwater sediment | Machu Picchu, PER | Mazencova et al., 2023 |
| OR291175 | Trepomonas sp.3 SCHOONER4 | Environment, freshwater | Freshwater sediment | Vancouver Island, CAN | Mazencova et al., 2023 |
| OR291169 | Trepomonas sp.4 POMPEJE | Environment, freshwater | Freshwater sediment | Pompeje, ITA | Mazencova et al., 2023 |
| OR291170 | Trepomonas sp.4 KAMERUN2 | Environment, freshwater | Freshwater sediment | CMR | Mazencova et al., 2023 |
| EF100197 | Uncultured eukaryote clone D1P02C | Environment, marine | oxygen-depleted intertidal marine sediment, upp2 cm | Greenland: Arctic | Stoeck,T. et al. 2006 |

| GenBank numb | Species | host | ENVIRONMENT | Land of origin | Referen |
|---|---|---|---|---|---|
| EF526748 | Uncultured marine eukar clone NA1 | Environment, marine | marine environment | Norway, Framvaren fjord | Behnke,A.et. al.. 2010 |
| EF526742 | Uncultured marine eukar clone NA1 | Environment, marine | marine environment | Norway, Framvaren fjord | Behnke,A.et. al.. 2010 |
| EF526744 | Uncultured marine eukar clone NA1 | Environment, marine | marine environment | Norway, Framvaren fjord | Behnke,A.et. al.. 2010 |
| OR291215 | Trepomonas agilis MIS2C | Environment, freshwater | Miskovice, CZE | Czech Republic | Mazancova,E. 2023 |
| U53120 | Trepomonas agilis | Environment, freshwater | Environment free living | | Cavalier-Smith,T. and Chao,E.E. 1996 |
| OR291216 | Trepomonas agilis SOOS4 | Environment, freshwater | Freshwater sediment, Soos, CZE | Czech Republic | Mazancova,E. 2023 |
| OR291196 | Trepomonas latecapitata BEAVER3 | Environment, freshwater | Freshwater sediment, Vancouver, CAN | Canada | Mazancova,E. 2023 |
| OR291197 | Trepomonas latecapitata PETRYB | Environment, freshwater | Freshwater sediment, Mladoˇsovice, CZE | Czech Republic | Mazancova,E. 2023 |
| OR291185 | Trepomonas steini CONGO | Environment, freshwater | Freshwater sediment, Brazzaville, COG | Congo | Mazancova,E. 2023 |
| OR291188 | Trepomonas steini GOLAKASAJ | Environment, freshwater | Freshwater sediment, Golakasaj, Rajasta´n, IND | IND | Mazancova,E. 2023 |
| OR291193 | Trepomonas steini KAIYKATAAN | Environment, freshwater | Freshwater sediment, Kaiykata, Darjeeling, IND | IND | Mazancova,E. 2023 |
| OR291189 | Trepomonas steini KAMERUN3 | Environment, freshwater | Freshwater sediment | CMR | Mazancova,E. 2023 |
| OR291186 | Trepomonas steini KUN | Environment, freshwater | Freshwater sediment, Orlick´e hory, | CZE | Mazancova,E. 2023 |
| OR291194 | Trepomonas steini LOUISE1 | Environment, freshwater | Freshwater sediment, Lake Louise | Alberta, CAN | Mazancova,E. 2023 |
| OR291190 | Trepomonas steini LUKIFOREST | Environment, freshwater | Freshwater sediment | Luki, COD | Mazancova,E. 2023 |
| OR291187 | Trepomonas steini LUH4 | Environment, freshwater | Freshwater sediment, river Vltava | CZE | Mazancova,E. 2023 |
| OR291192 | Trepomonas steini MOSSE1 | Environment, freshwater | Freshwater sediment | CAF | Mazancova,E. 2023 |
| OR291205 | Trepomonas rotans KIEL9 | Environment, freshwater | Freshwater sediment, | Behrensdorf, DEU | Mazancova,E. 2023 |
| OR291204 | Trepomonas rotans MALSE | Environment, freshwater | Freshwater sediment, ˇ | Ceske' Budˇejovice, CZE | Mazancova,E. 2023 |
| OR291225 | Trepomonas rotans vT2 | Environment, freshwater | Freshwater sediment, pond | Velký Tisý, CZE | Mazancova,E. 2023 |
| OR291198 | Trepomonas rotans BOL2 | Environment, freshwater | Freshwater sediment, | NP Eduardo Avaroa, BOL | Mazancova,E. 2023 |
| OR291208 | Gyromonas ambulans LUH2N | Environment, freshwater | Freshwater sediment, Vltava river alluvial plain, | CZE | Mazancova,E. 2023 |
| OR291206 | Gyromonas ambulans GRUBER2 | Environment, freshwater | Freshwater sediment, | Grüber, Brunnen, DEU | Mazancova,E. 2023 |
| OR291207 | Gyromonas ambulans SPINDL2 | Environment, freshwater | Freshwater sediment, | Sˇpindlerův Mlýn, CZE | Mazancova,E. 2023 |
| OR291218 | Hexamita sp. AYEN | Environment, freshwater | Freshwater sediment | region Aysen, CHL | Mazancova,E. 2023 |
| HF568859 | uncultured eukaryote | Environment, freshwater | groundwater sample | USA:Colorado, Rifle | Holmes,D.E.2013 |
| HF568849 | uncultured Hexamita RifleRDL16 | Environment, freshwater | groundwater sample | USA:Colorado, Rifle | Holmes,D.E.2013 |
| KJ566522 | Uncultured euryarchaeote SM_S_2 | Environment, freshwater | cold freshwater spring | Germany? | Moissl-Eichinger,C. 2014 |
| HF568847 | uncultured Trepomonas RifleRDL15 | Environment, freshwater | groundwater sample | USA:Colorado, Rifle | Holmes,D.E.2013 |
| OR291211 | Hexamita sp. BOTAN1 | Environment, freshwater | Freshwater sediment, Praha , CZE | | Mazancova,E. 2023 |
| OR291224 | Hexamita sp. KAMERUN4 | Environment, freshwater | Freshwater sediment, CMR | | Mazancova,E. 2023 |
| OR291219 | Hexamita sp. KLOSTERSEE | Environment, freshwater | Freshwater sediment, Klostersee, DEU | | Mazencova et al., 2023 |
| OR291220 | Hexamita sp. NORSKO | Environment, Brackish/Ma | NOR | | Mazencova et al., 2023 |
| AY916608 | uncultured eukaryote Zeuk103 | Environment, freshwater | sulfide-rich Zodletone spring | USA: Oklahoma | Luo,Q., 2005 |
| AY916607 | uncultured eukaryote Zeuk104 | Environment, freshwater | sulfide-rich Zodletone spring | USA: Oklahoma | Luo,Q., 2005 |
| OR291221 | Hexamita sp. SEB5 | Environment, freshwater | pond Sˇeber´ak, Praha, CZE | | Mazancova,E. 2023 |
| OR291212 | Hexamita sp. ZOO1 | Environment, freshwater | ZOO Praha, CZE | | Mazancova,E. 2023 |
| OR291223 | Trimitus sp. VAV2A | Environment, freshwater | Vejvanov-Pajzov, CZE | | Mazancova,E. 2023 |
| OR291213 | Trepomonas sp. ESMI3 | *Eudicella smithi bertheranc* | Larvae of flower beetle, originating from East Africa | | Mazancova,E. 2023 |
| OR291214 | Trepomonas sp. EUDIA3 | *Eudicella* sp., larva | Larvae of flower beetle | | Mazancova,E. 2023 |
| OR291209 | Hexamita sp. PI1kl3 | *Haemopis sanguisuga* | intestine horse leech, pond | Prag, CZE | Mazancova,E. 2023 |
| OR291210 | Hexamita sp. PI1kl5 | *Haemopis sanguisuga* | intestine horse leech, pond | Prag, CZE | Mazancova,E. 2023 |
| OR291217 | Hexamitinae sp. CERAT3 | *Ceratophrys ornata* . | fecal sample, Argentine horned frog | ? | Mazancova,E. 2023 |
| OR291222 | Hexamitinae sp. SCHULTZ2 | *Cetoniinae* sp.larva | intestine Beetle larvae | ? | Mazancova,E. 2023 |
| OR291226 | Hexamitinae sp. TIPLICE | *Tipula* sp., larva | intestine Cranefly larvae | ? | Mazancova,E. 2023 |
