## Supplementary Figure 2 for "First report of *Hexamita nelsoni* in blue mussel (*Mytilus edulis*): morphology, phylogeny, and host-parasite interaction"

Supplementary Figure 1: phasecontrast *Hexamita nelsoni* from cell culture (LK22-14) of *Hexamita nelsoni* vizualising differently shaped single trophozoites (A-I) and cystlike cells (J-L).

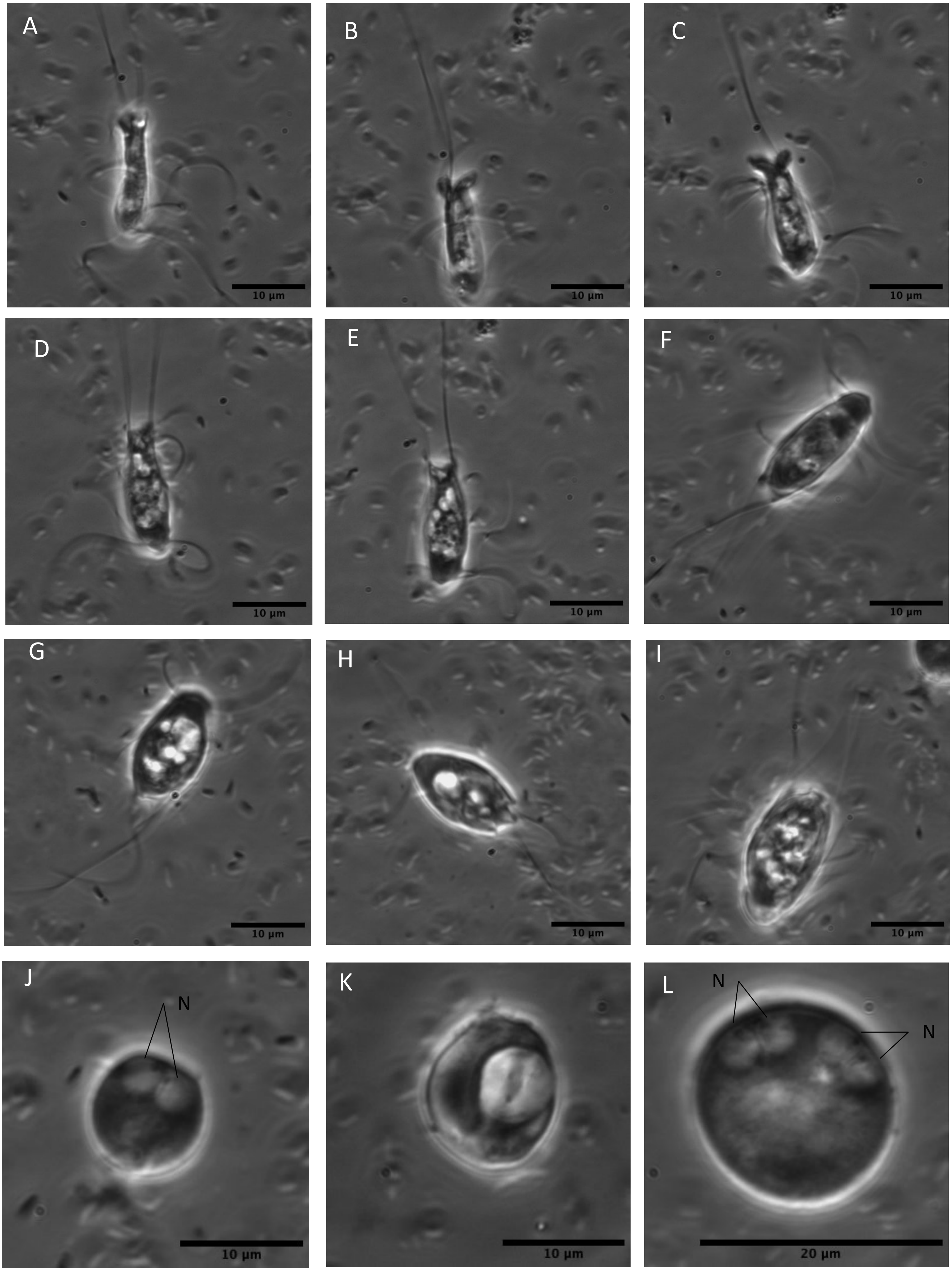
