## Supplementary Figure 3 for "First report of *Hexamita nelsoni* in blue mussel (*Mytilus edulis*): morphology, phylogeny, and host-parasite interaction"

Supplementary Figure 2: DIC (A,D,G )and phasecontrast (B-C, E-F, H-L) from cell culture (LK22-14) of *Hexamita nelsoni* vizualising large trophozoites, enclosing double cells enclosed within one membrane.

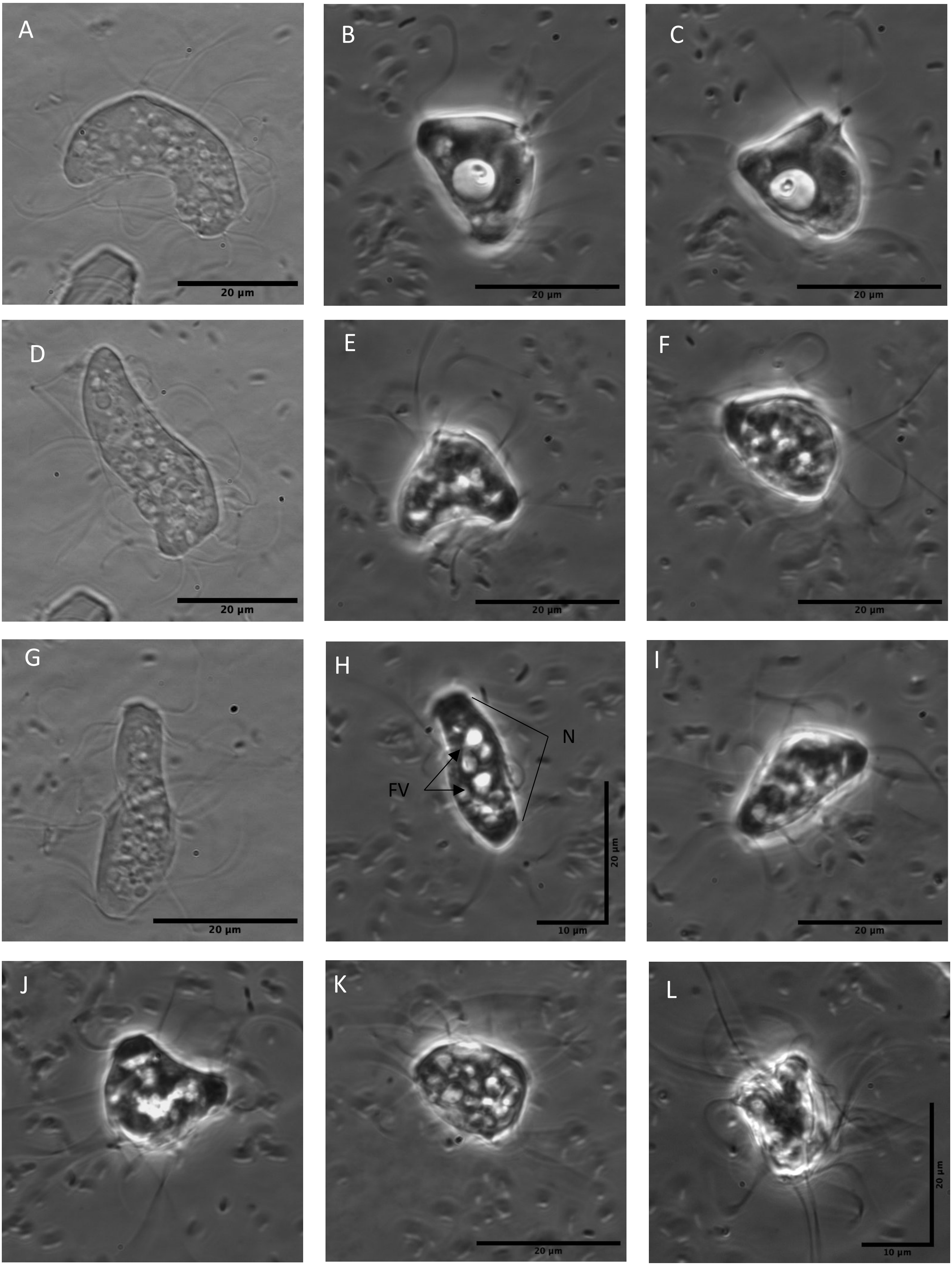
